# Intestinal norovirus binding patterns in non-secretor individuals

**DOI:** 10.1101/2022.05.26.493674

**Authors:** Georges Tarris, Marie Estienney, Philippe Daval-Frérot, Anne-Cécile Lariotte, Damien Aubignat, Karine Sé, Christophe Michiels, Laurent Martin, Alexis de Rougemont, Gaël Belliot

## Abstract

Human norovirus (HuNoV) infection is associated with active FUT2 status, which characterizes the secretor phenotype. However, non-secretor individuals are also affected by HuNoV infection although in a lesser proportion. Here, we study GII.3, GII.4 and GII.17 HuNoV interactions in non-secretor individuals using baculovirus-expressed virus-like particles (VLPs). Only GII.4 HuNoV specifically interacted with non-secretor saliva. Competition experiments using HBGA-specific mAbs demonstrate that GII.4 VLPs recognized the Lewis a antigen (Le^a^). We also analyzed HuNoV VLP interactions on duodenum tissue blocks from healthy non-secretor individuals. VLP binding was observed for the three HuNoV genotypes in 10 of the 13 individuals, and competition experiments demonstrated that VLP recognition was driven by interaction with the Le^a^ antigen. In 3 individuals, binding was restricted to either GII.4 alone or GII.3 and GII.17. One patient did not display VLP binding for any of the three genotypes.

Finally, we performed a VLP binding assay on proximal and distal colon tissue blocks from a non-secretor patient with Crohn’s disease. VLP binding to inflammatory tissues was genotype-specific since GII.4 and GII.17 VLPs were able to interact with regenerative mucosa whereas GII.3 VLP was not. Binding of GII.4 and GII.17 HuNoV VLPs was linked to Le^a^ in regenerative mucosae from the proximal and distal colon. Overall, our data clearly showed that Le^a^ has a pivotal role in the recognition of HuNoV in non-secretors. We also showed that Le^a^ is expressed in inflammatory/regenerative tissues and that it can interact with HuNoV in secretor and non-secretor individuals. The physiological and immunological consequences of such interactions in non-secretors has yet to be elucidated.

**IMPORTANCE:** Human norovirus (HuNoV) is the main etiological agent of viral gastroenteritis in all age classes. HuNoV infection mainly affects secretor individuals, who are characterized by the presence of the ABO(H) and Lewis histo-blood group antigens at the surface of the small intestine. Non-secretor individuals, who only express Lewis antigens (Le), are less susceptible to HuNoV infection. Here we study the interaction of three frequently encountered HuNoV genotypes (GII.3, GII.4 and GII.17) in non-secretor individual using baculovirus-expressed viral particles. Preliminary saliva binding assays showed that only GII.4 interacted with non-secretor saliva via the Le^a^ antigen.

Surprisingly, in the binding assays on duodenal tissue blocks, the three genotypes interacted with non-secretor enterocytes via Le^a^. This suggests that HBGA status in the saliva does not necessarily reflect interactions in the intestines and, secondly, that Le^a^ plays a pivotal role in HuNoV attachment in non-secretors. Similarly, Le^a^ was involved in the recognition of GII.4 and GII.17 HuNoV particles by inflammatory colon tissue from a non-secretor Crohn’s disease patient. The molecular implications of HuNoV binding in non-secretors remains to be elucidated in physiological and pathological conditions encountered in other intestinal diseases.

## INTRODUCTION

Human norovirus (HuNoV) is the main etiological agent of gastroenteritis in all age classes worldwide (1, 2). After the discovery of HuNoVs in 1972, the lack of cell culture system hampered its study until 2016, when the first robust system for norovirus cultivation was conceived using human intestinal enteroids (HIE) (3-5). Previous studies showed that HuNoV replication only occurs in organoids derived from the small intestine, so colonoids cannot be used for HuNoV replication (5-7). Similarly, histological analysis of tissues derived from immunocompromised patients with chronic HuNoV infection revealed that HuNoV replicates in jejunal enterocytes and, more surprisingly, in enteroendocrine cells (8). The use of HIE confirmed the pivotal role of the *FUT2* gene during HuNoV replication, while no binding was observed with HIEs derived from non-secretor individuals, except for the GII.3 genotype (6, 9). Before the use of HIEs, the *FUT2* gene was already associated with susceptibility to HuNoV infection. The *FUT2* gene encodes α1,2 fucosyltransferase, which regulates the expression of histo-blood group antigens (HBGAs) in various mucosa, including in the small intestine, and in secretions like saliva (10). HBGAs are displayed at the surface of enterocytes and are the only known ligand involved in the recognition of HuNoV (10). The active *FUT2* gene defines the secretor genotype, which is 80% predominant in the European population. For the other 20%, who are known as non-secretors, the *FUT2* gene is inactivated by nonsense mutations, among which G428A is the most frequently encountered within the Caucasian population (11, 12). The mutation of both alleles characterizes the recessive non-secretor phenotype, whose main feature is the absence of *FUT2*-dependent HBGAs (i.e., A, B, H antigens) in saliva and at the surface of enterocytes (13). The expression of Lewis antigens is driven by the *FUT3* gene and is conditioned by the status of the *FUT2* gene. The *FUT3* gene is inactive in nearly 10% of the European population (10, 13). In secretor individuals, Lewis b (Le^b^) and Lewis y (Le^y^) are expressed in saliva and mucosa whereas Lewis a (Le^a^) and Lewis x (Le^x^) antigens are only expressed in non-secretor individuals (13, 14). Therefore, secretor and non-secretor individuals are in most cases Le^a-^Le^b+^ and Le^a+^Le^b-^, respectively. The role of HBGAs during HuNoV infection has been largely explored in healthy patients, and to a lesser extent in patients diagnosed with inflammatory bowel disease (IBD) (15). IBD includes several progressive digestive diseases including Crohn’s disease and ulcerative colitis. Ulcerative colitis only affects the colon and rectum while Crohn’s disease can be extended to the entire digestive tract with possible extra-intestinal manifestations (i.e., ocular, articular or cutaneous) (16). The course of IBD is characterized by alternating episodes of remission and flare up (17). The symptoms (i.e. abdominal pain, fatigue, diarrhea, weight loss) are similar for both diseases (16). Although the onset of IBD remains unexplained, the triggering factors appear to involve a combination of immunological, environmental, genetic and microbiological factors (18, 19). The intestinal lumen of IBD patients is characterized by simplified glycan structures, which might favor accidental infection by enteric pathogens and increase the risk of inflammation (20, 21). During flare ups, marked structural alterations are seen in the inflamed mucosae (16). Blood group ABO(H) antigens are usually not displayed at the surface of inflammatory and regenerative colonic and rectal mucosa, whereas a strong expression of Sialyl-Le^a^ (sLe^a^) (also known as CA19.9) and to a lesser extent Sialyl-Le^x^ (sLe^x^) (also known as CD15s) is observed (15, 22-24). The absence of sLe^a^ antigen expression on quiescent mucosa contrasts with its reversible expression in regenerative mucosa and its persistent expression in dysplasia (22). The putative link between IBD and HBGA expression has not fully been demonstrated yet. According to genetic studies conducted in European populations, non-secretor individuals have an increased risk of Crohn’s disease-associated ileitis (25-28). The role of viral agents as a cause or consequence of IBD and the related dysbiosis remain largely debated, as exemplified for bacteriophages (29). Since HBGAs and *FUT2* status are strongly related to HuNoV infections, several epidemiological studies were performed to determine whether the occurrence of HuNoV infections is increased in IBD patients. The results remain contrasted: initial studies based upon small cohorts showed no association between enteric viral infections and IBD while three recent larger-scale studies demonstrated that Crohn’s and colitis patients who suffered from flare-ups had a significantly higher prevalence of HuNoV infection (30-34). The weaker interactions between glycans and enteric pathogens in non-secretor subjects might play a role in immune dysregulation and IBD pathogenesis (20). Similar conclusions were reached for ulcerative colitis where *FUT3* polymorphisms were associated with the disease in Chinese populations (35). Previous studies showed that sLe^a^ and sLe^x^ antigens were strongly expressed at the surface of regenerative mucosa in secretor patients with refractory Crohn’s disease or ulcerative colitis (15). Additionally, Le^a^ and to a lesser extent Le^x^ antigens were responsible for the specific attachment of GII.4 HuNoV in inflamed tissues. The biological significance of HuNoV interactions with inflamed tissues in IBD remains unexplained in secretor and non-secretor individuals.

Here, we characterize GII.4, GII.3 and GII.17 HuNoV binding patterns and HuNoV-HBGA interactions in the saliva and healthy duodenal tissues of non-secretor individuals. We then characterized carbohydrates that are involved in HuNoV binding. We finally analyzed the binding patterns of HuNoV and its interactions with HBGA in the inflammatory tissues of a non-secretor patient diagnosed with Crohn’s disease.

## RESULTS

### Norovirus VLP binding profiles in non-secretor saliva

Thirty-four genotyped saliva samples from non-secretor individuals (Le^a+^ Le^b-^) were used for binding assays using GII.3, GII.4 and GII.17 VLPs (Figure 1). GII.4 VLP detection was abolished following sodium periodate treatment, which suggests that carbohydrates were involved in VLP recognition (data not shown). GII.4 Osaka variant have been previously documented as binding to non-secretor saliva (36, 37). GII.4 binding was therefore used as a positive control for the experiments. For the cut-off, the limit was arbitrarily fixed at 0.2 OD_450nm_. Binding amplitudes to non-secretor saliva were significantly lower than in secretor saliva, as previously described (36). GII.4 VLPs were detected in 15 out of 34 salivary samples, with OD_450nm_ values ranging from 0.211 to 0.69 (mean OD for the positive samples: 0.41±0.17). GII.17 Kawasaki 308 VLP were detected in 2 out of 34 salivary samples, with OD_450nm_ values higher than 0.3. It is worth noting that, along with GII.4 VLPs, the positive salivary samples for GII.17 VLPs were among the most positive (0.64 for sample 5 and 0.62 for sample 22). The GII.3 SW4 strain did not bind to any of the non-secretor saliva, confirming previous results (37).

**Figure 1:**
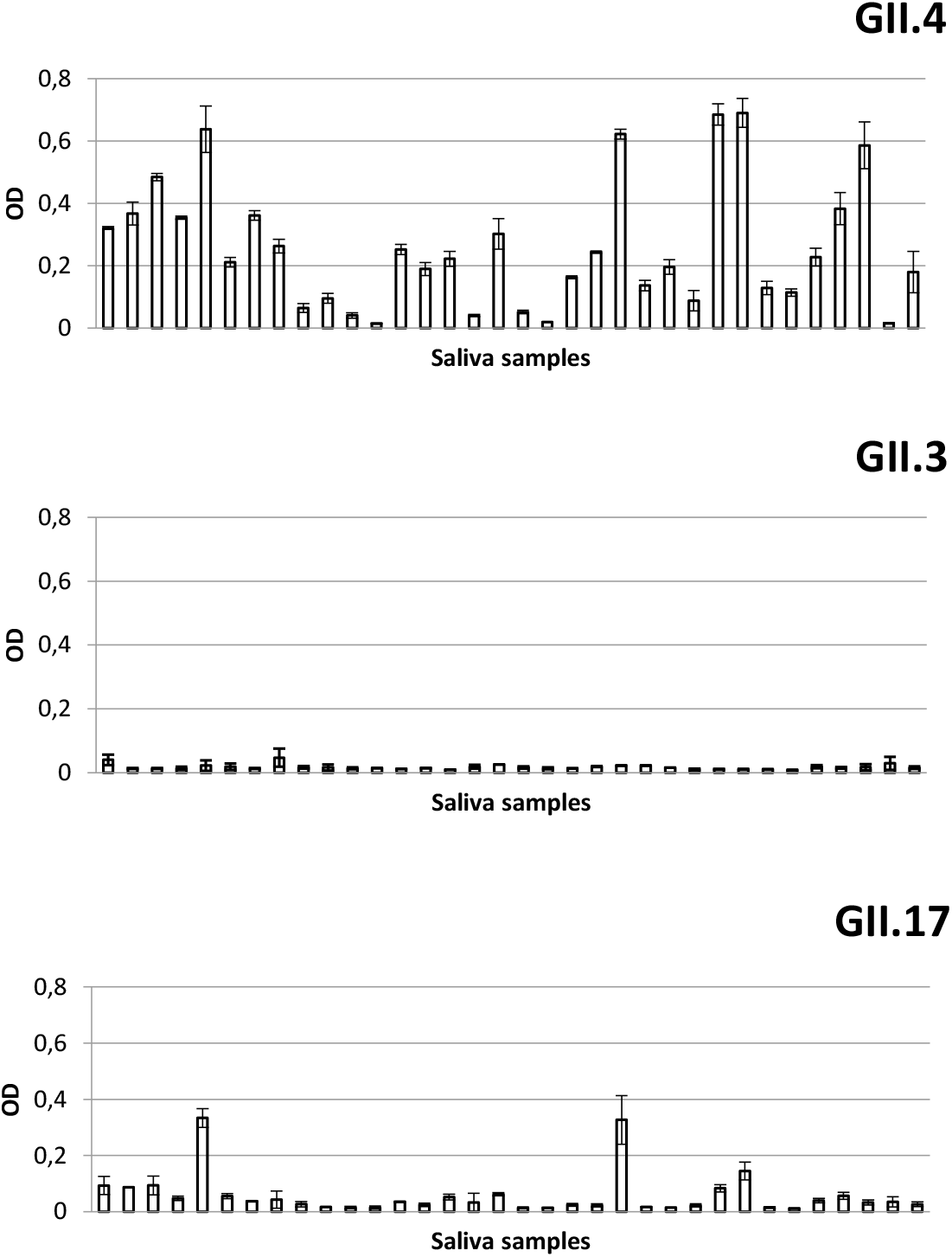
Saliva binding assays on non-secretor saliva. The binding of cesium chloride-purified VLP was measured by ELISA in duplicate for each saliva sample, and the mean values are shown on the graph. The VLP genotype is indicated on the top right corner of each histogram. The number of the sample is indicated on the horizontal axis, and OD_450nm_ values are indicated on the vertical axis of each histogram.

### Role of Lewis antigens into VLP attachment to non-secretor saliva

To determine the role of HBGA in GII.4 binding in saliva, competition experiments were performed using four of the saliva samples from the cohort with a sufficient volume (salivary samples 2, 3, 5 and 6) (Figure 2). Unfortunately, the OD_450nm_ values for GII.17 VLP were too low to attempt a similar competition experiment. For each sample, the coated saliva was first incubated with increasing amounts of Lewis-specific purified monoclonal antibodies (mAbs). For the Le^b^ and Le^x^ antigens, increasing amounts of specific mAbs did not inhibit VLP binding to the saliva, which suggested that the Le^b^ and Le^x^ antigens were not involved in GII.4 VLP attachment. Inversely, a marked inhibition of the GII.4 VLP attachment was observed in the four saliva samples following preincubation of increasing amounts of Le^a^-specific mAbs, demonstrating that GII.4 VLP binding to non-secretor saliva involves Le^a^ antigen. In the next set of experiments, immunohistochemical analyses were conducted using tissue samples derived from healthy non-secretor individuals to confirm the role of the Le^a^ ligand in HuNoV attachment.

**Figure 2:**
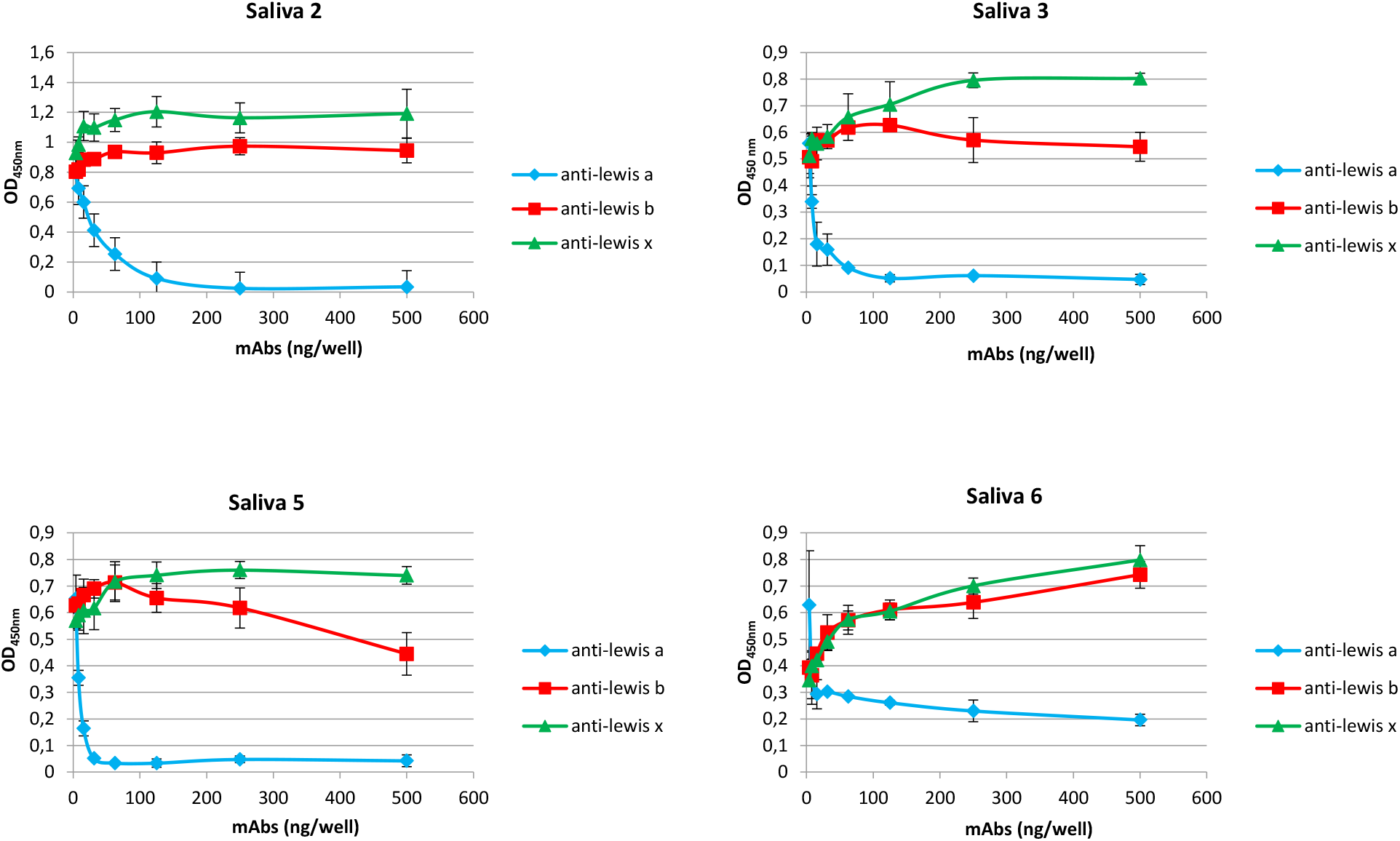
Competition experiments using GII.4 VLP in saliva binding assays. Each experiment was conducted in triplicate for each saliva sample, and the mean values are shown on the graph. Vertical bars represent standard deviation. The saliva sample is indicated above each graph. On each graph, antibodies are color-coded (anti-Le^a^ (blue), anti-Le^b^ (red) and anti-Le^x^ (green)) and the legend is on the right side of the graph. The OD_450nm_ values and mAbs quantity used for the experiments are indicated on the vertical axis and the horizontal axis, respectively.

### HuNoV interactions on healthy duodenum from non-secretor individuals

Thirteen healthy duodenum samples (Patients 1NS to 13NS) were selected for the study (mean age: 53 years, sex ratio: 11F/2M). For the blood groups, 5 patients were type O, 7 patients were type A and 1 patient was type B (Table 1). None of the patients displayed ABH(O) antigens at the surface of duodenal villi (data not shown). For patients 1NS through 9NS, GII.3, GII.4 and GII.17 VLP binding was detected on the surface of duodenal villi (Figure 3). For patient 10NS, VLP binding was restricted to GII.4 VLP at the surface of duodenal enterocytes (Figure 3). In patients 11NS and 12NS, a strong binding signal was observed for GII.3 and GII.17 VLPs but no binding was observed for GII.4 VLPs. Patient 13NS did not display VLP binding for any of the three genotypes. As for HBGA detection, the duodenal biopsies of patients 1NS to 9 NS strongly expressed Le^a^ in villous goblet and epithelial cells, contrary to the biopsies of patients 10NS, 11NS and 13NS which showed a weak expression of the Le^a^ antigen (Figure 4). The Le^x^ antigen was constitutively expressed in all cases. The expression of Le^y^ was restricted to the base of the crypts in all cases and was absent from the apical side of the duodenum. Overall, our data suggest the correlation between HuNoV binding and Lewis antigen expression in the duodenum in non-secretor individuals in the absence of ABH(O) expression, as demonstrated in salivary samples. To go further, we attempted to identify which Lewis antigen was involved in the interactions between VLPs and healthy duodenal samples from non-secretor individuals.

**Table 1:**
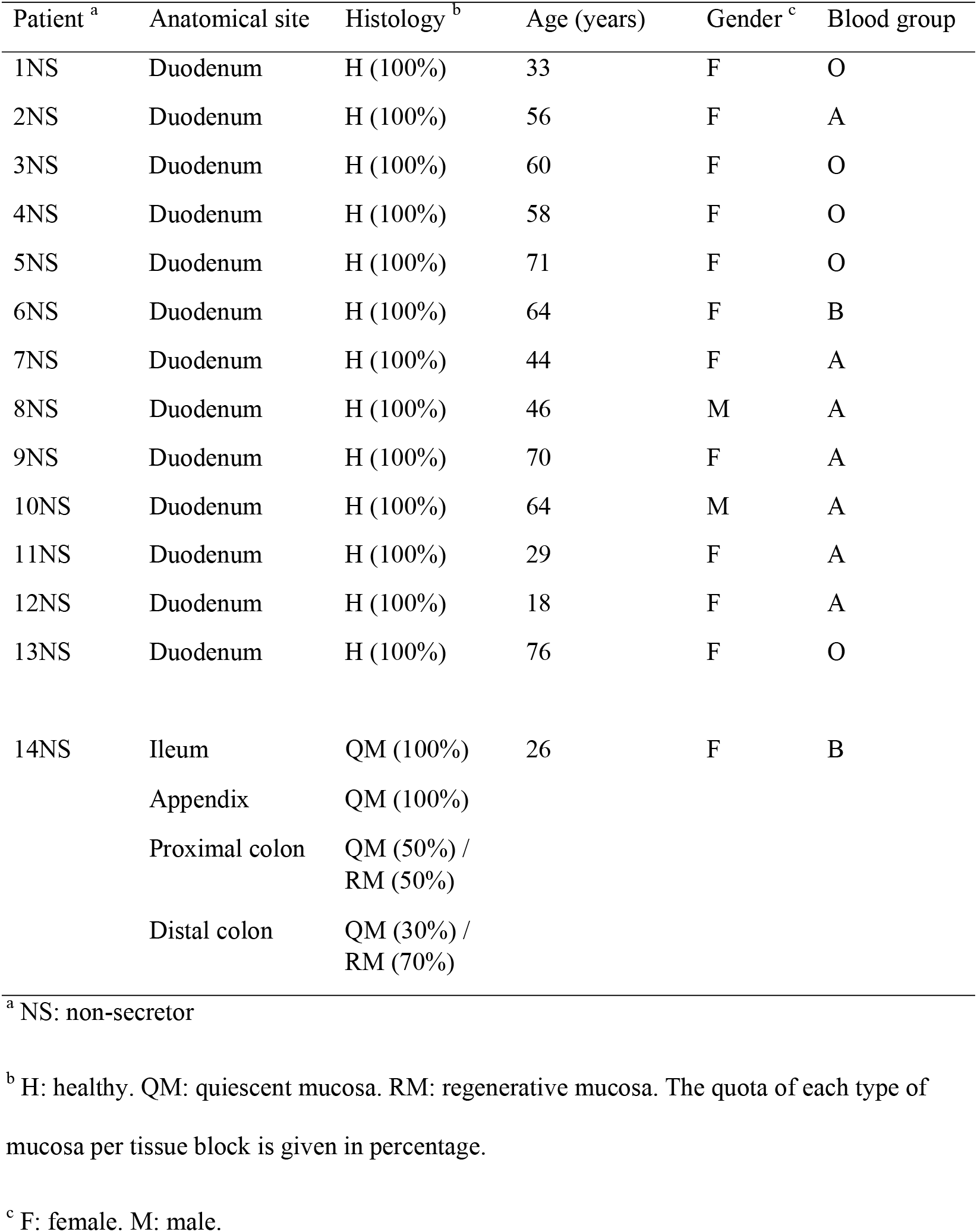
Cohort used for the study

| Patient <sup>a</sup> | Anatomical site | Histology <sup>b</sup> | Age (years) | Gender <sup>c</sup> | Blood group |
| --- | --- | --- | --- | --- | --- |
| 1NS | Duodenum | H (100%) | 33 | F | O |
| 2NS | Duodenum | H (100%) | 56 | F | A |
| 3NS | Duodenum | H (100%) | 60 | F | O |
| 4NS | Duodenum | H (100%) | 58 | F | O |
| 5NS | Duodenum | H (100%) | 71 | F | O |
| 6NS | Duodenum | H (100%) | 64 | F | B |
| 7NS | Duodenum | H (100%) | 44 | F | A |
| 8NS | Duodenum | H (100%) | 46 | M | A |
| 9NS | Duodenum | H (100%) | 70 | F | A |
| 10NS | Duodenum | H (100%) | 64 | M | A |
| 11NS | Duodenum | H (100%) | 29 | F | A |
| 12NS | Duodenum | H (100%) | 18 | F | A |
| 13NS | Duodenum | H (100%) | 76 | F | O |
| 14NS | Ileum | QM (100%) | 26 | F | B |
|  | Appendix | QM (100%) |  |  |  |
|  | Proximal colon | QM (50%) /<br>RM (50%) |  |  |  |
|  | Distal colon | QM (30%) /<br>RM (70%) |  |  |  |

<sup>a</sup> NS: non-secretor
<sup>b</sup> H: healthy. QM: quiescent mucosa. RM: regenerative mucosa. The quota of each type of mucosa per tissue block is given in percentage.
<sup>c</sup> F: female. M: male.

**Figure 3:**
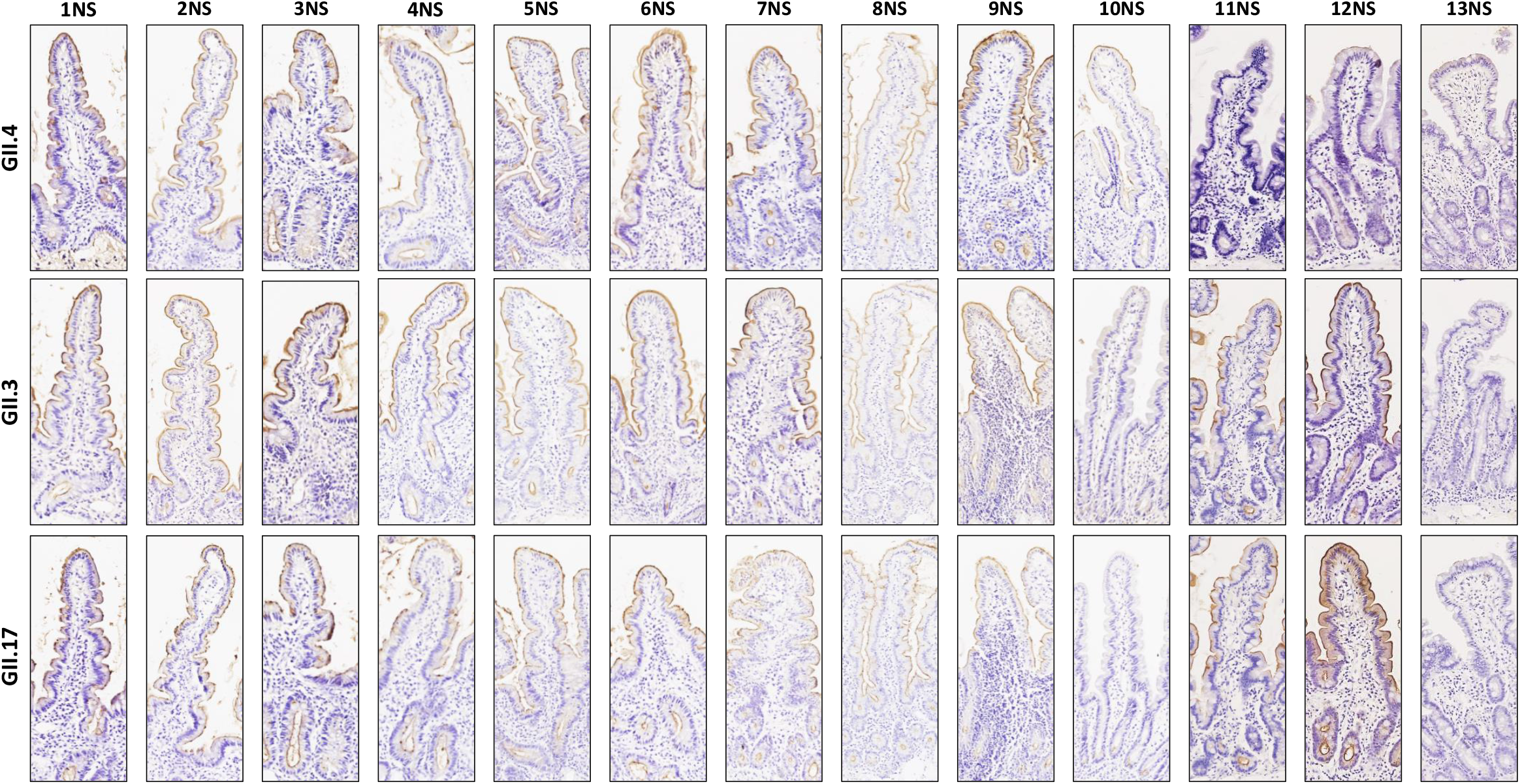
GII.4, GII.3 and GII.17 binding in healthy non-secretor duodenal samples. Incubated VLP are indicated on the left of each line. Duodenal samples 1NS to 13NS are indicated on the top of each row. Positive VLP binding is characterized by brown staining. The panels of this figure and the following figure are shown at magnification x200 unless indicated otherwise.

**Figure 4:**
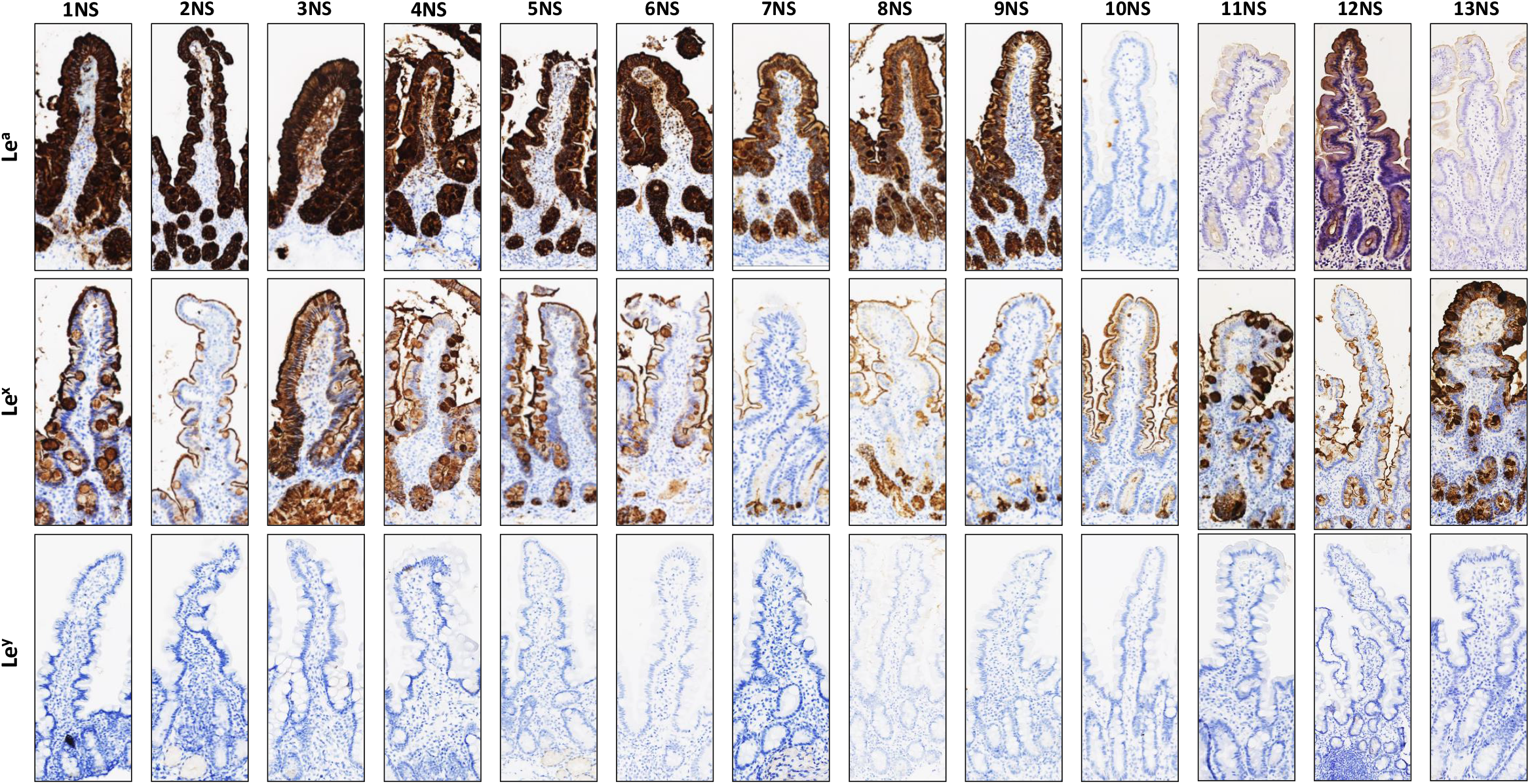
Lewis antigen detection in healthy non-secretor duodenal samples. The detected Lewis antigens are indicated to the left of each line. Duodenal samples 1NS to 13NS are indicated on top of each row. Positive HBGA detection is shown by brown staining.

### Characterization of GII.4, GII.3 and GII.17 attachment to healthy non-secretor duodenum

In the absence of expression of ABH(O) blood group antigens, the occurrence of VLP binding at the surface of duodenal villi might be explained by the expression of other HBGAs such as Lewis antigens. To verify this hypothesis, duodenal biopsies from patients 3NS, 11NS and 12NS were selected for competition experiments using Lewis-specific mAbs. Limited sample sizes for the three biopsies forced us to perform the experiment on one tissue for each genotype. Slides were preincubated with Le^a^ and either Le^b^- or Le^x^-specific mAbs, combined or alone. Slides were then incubated with GII.4 VLP on sample 3NS, GII.3 VLP on sample 12NS and GII.17 VLP on sample 11NS (Figure 5). Preincubation of Le^a^-specific mAbs totally suppressed VLP binding for the three genotypes, but preincubation of Le^x^-(GII.4) and Le^b^-specific mAbs (GII.3 and GII.17), did not. Our experiments demonstrated that the Le^a^ antigen plays a pivotal role in GII.4, GII.3 and GII.17 HuNoV VLP binding in the duodenal villi of non-secretor individuals, in the absence of ABH(O) expression.

**Figure 5:**
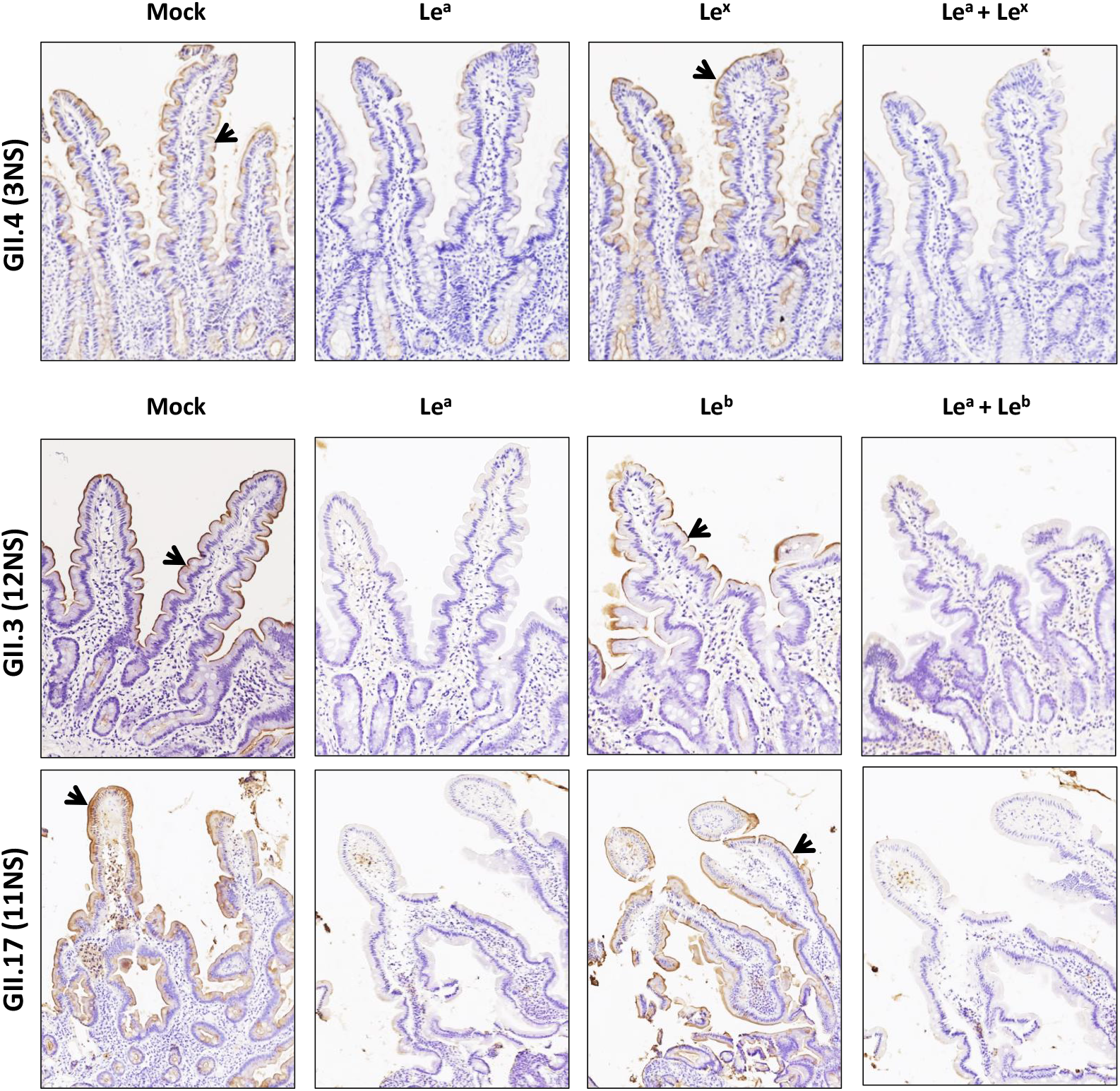
VLP competition experiments using healthy non-secretor duodenal samples from patients 3NS, 12NS and 11NS. Preincubation of mAbs and control are indicated on top of each row. The genotype of the VLP used for the assay is indicated to the left of each line. The sample used for the experiments is indicated in parentheses. Positive VLP detection is indicated by brown staining and with arrows.

### HuNoV interactions with inflammatory tissues from a non-secretor Crohn’s disease patient

A 33-year-old female patient with blood group B and suffering from refractory Crohn’s disease (Patient 14NS) was selected following the retrospective screening of our cohort (Table 1). The patient had undergone subtotal colectomy, after which sampling was performed in the Pathology Department of the Dijon University Hospital (Dijon, France). We observed quiescent mucosa and regenerative mucosa from the proximal and distal colonic tissues, as well as quiescent mucosa from appendiceal and ileal tissues (Table 1). Our experiments aimed to determine GII.4, GII.3 and GII.17 HuNoV VLP binding profiles in both quiescent and inflamed mucosa for a non-secretor individual. The following objective was to determine whether the observed binding capacity was the same in salivary samples and histological tissues.

Histological analysis revealed GII.3, GII.4 and GII.17 VLP binding to the quiescent mucosa of the ileum, in contrast to the absence of GII.3 VLP binding observed in healthy colonic tissues (Figure 6). For GII.4 VLPs, binding was detected in the bases of crypts from quiescent mucosa of the distal colon. The appendiceal quiescent mucosa showed GII.4 and GII.17 VLP detection in the upper crypt compartments. As for HBGA expression, none of the tissue samples expressed B antigens (Figure 7). Le^a^ and Le^x^ antigens were expressed in epithelial cells and goblet cells of the ileum, while sLe^a^ expression was restricted to the villous compartment. For the proximal colon and appendix, the quiescent mucosa displayed Le^a^, Le^x^, and to a lesser extent sLe^a^ antigens in goblet cells. The Le^a^, Le^x^ and sLe^a^ antigens were weakly expressed in the distal colon. For all samples, sLe^x^ and Le^y^ antigens were weakly expressed in the crypt bases.

**Figure 6:**
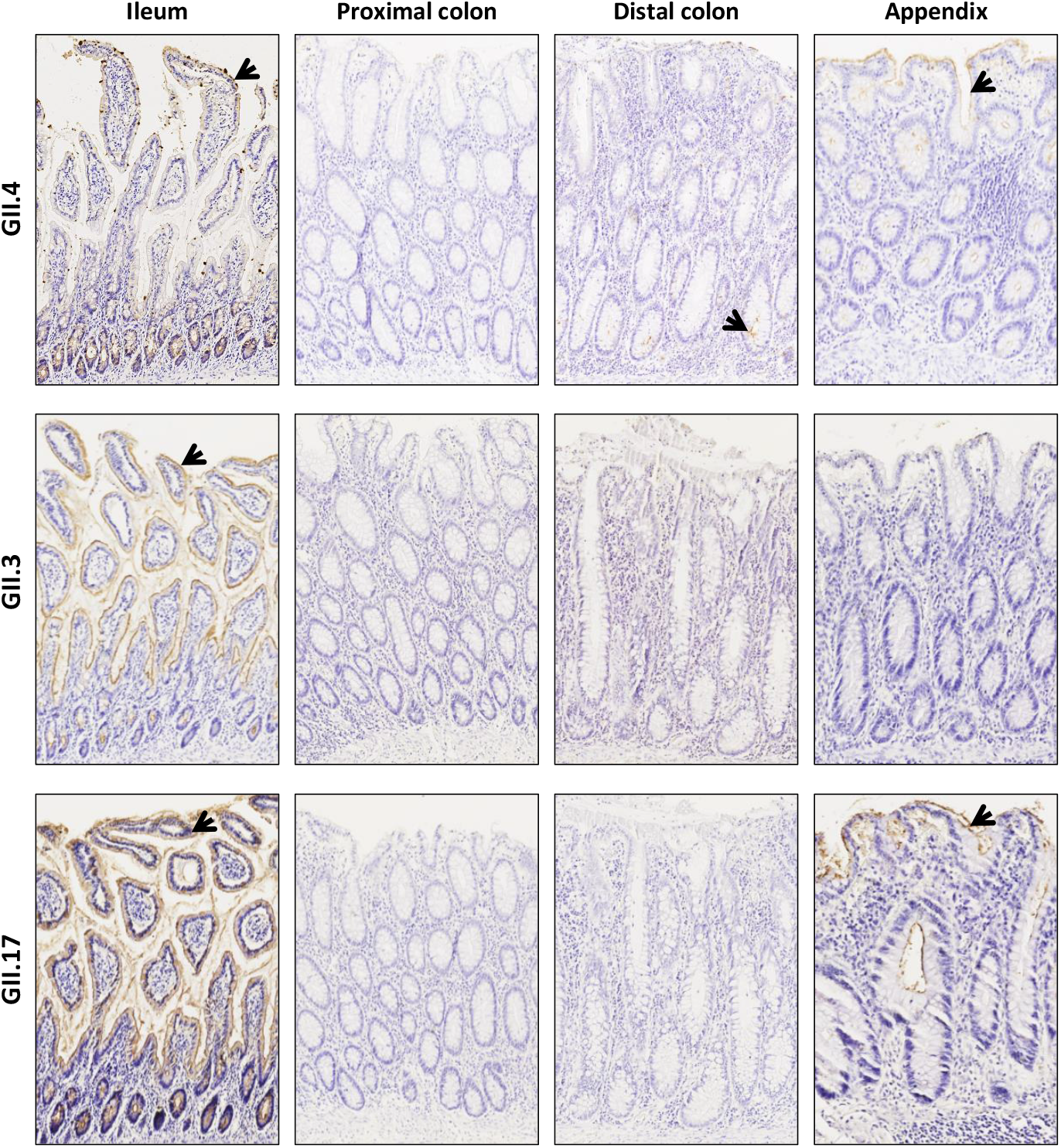
VLP detection in quiescent samples of the ileum, proximal colon, distal colon, and appendix from non-secretor Crohn’s disease patient 14NS. The anatomical site of the sample is indicated on top of each row. Incubated VLP are indicated at the left of each line. Positive VLP detection is shown by brown staining and with arrows.

**Figure 7:**
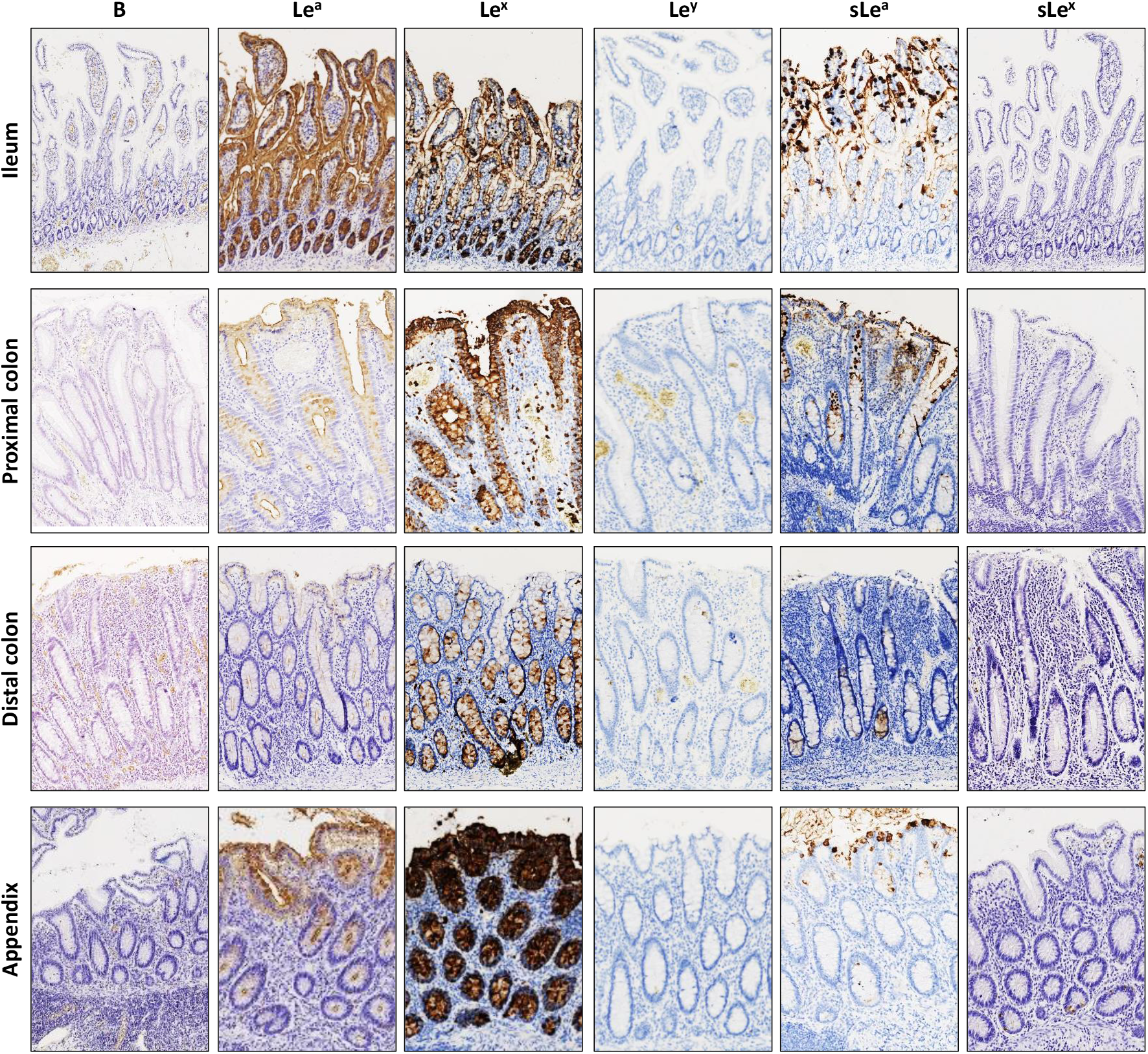
Blood group B, Lewis antigens and sialylated Lewis antigens detection in quiescent samples of the ileum, proximal colon, distal colon, and appendix from non-secretor Crohn’s disease patient 14NS. The anatomical origin of each tissue sample is indicated to the left of each line. The detected HBGA are indicated on top of each row. Positive HBGA detection is shown by brown staining.

Analysis of the regenerative mucosa of the proximal and distal colon revealed GII.4 and GII.17 VLP binding, but there was an absence of GII.3 VLP binding (Figure 8). Concerning HBGA detection in the regenerative mucosa of the distal and proximal colon, neither expressed the B antigen. However Le^a^, Le^x^, sLe^a^ and sLe^x^ were strongly expressed at the surface of regenerative epithelial cells (Figure 9). The Le^y^ antigen was not expressed in the regenerative mucosa of the proximal or distal colon. At this stage, our experiments indicate that VLP binding to inflammatory tissues is genotype-specific. Additionally, we observed that HuNoV VLP binding only occurred in inflammatory and regenerative tissue samples from this non-secretor individual suffering from Crohn’s disease, while there was no interaction between HuNoV VLPs and quiescent mucosa, as previously described in secretor individuals (16).

**Figure 8:**
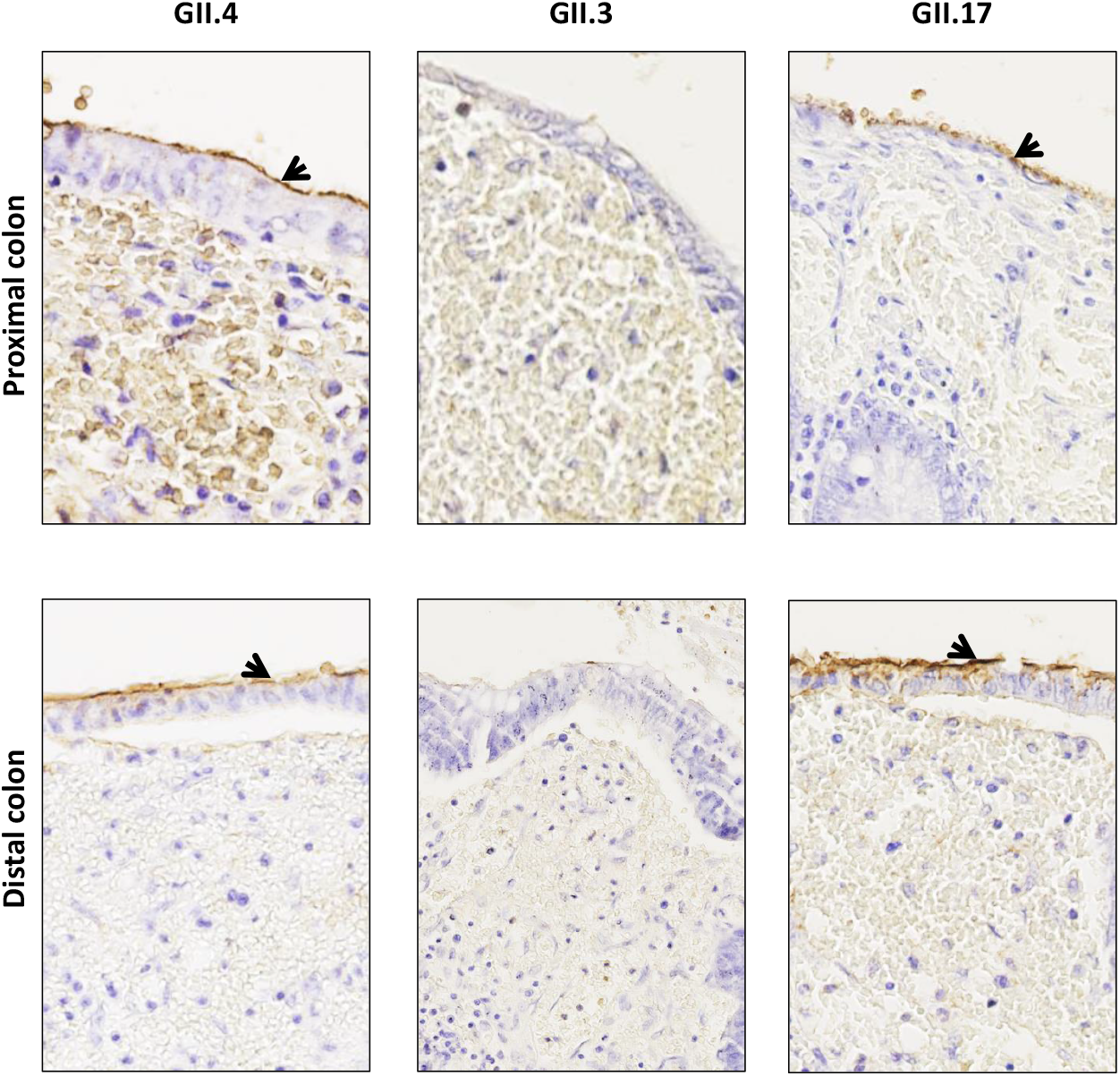
VLP detection in regenerative mucosa of samples of proximal and distal colon from non-secretor Crohn’s disease patient 14NS (magnification x400). The anatomical site of each sample are indicated on the left side of the panel. The VLP genotype is indicated above the panels (row). Positive VLP detection is indicated by brown staining and pointed by arrows.

**Figure 9:**
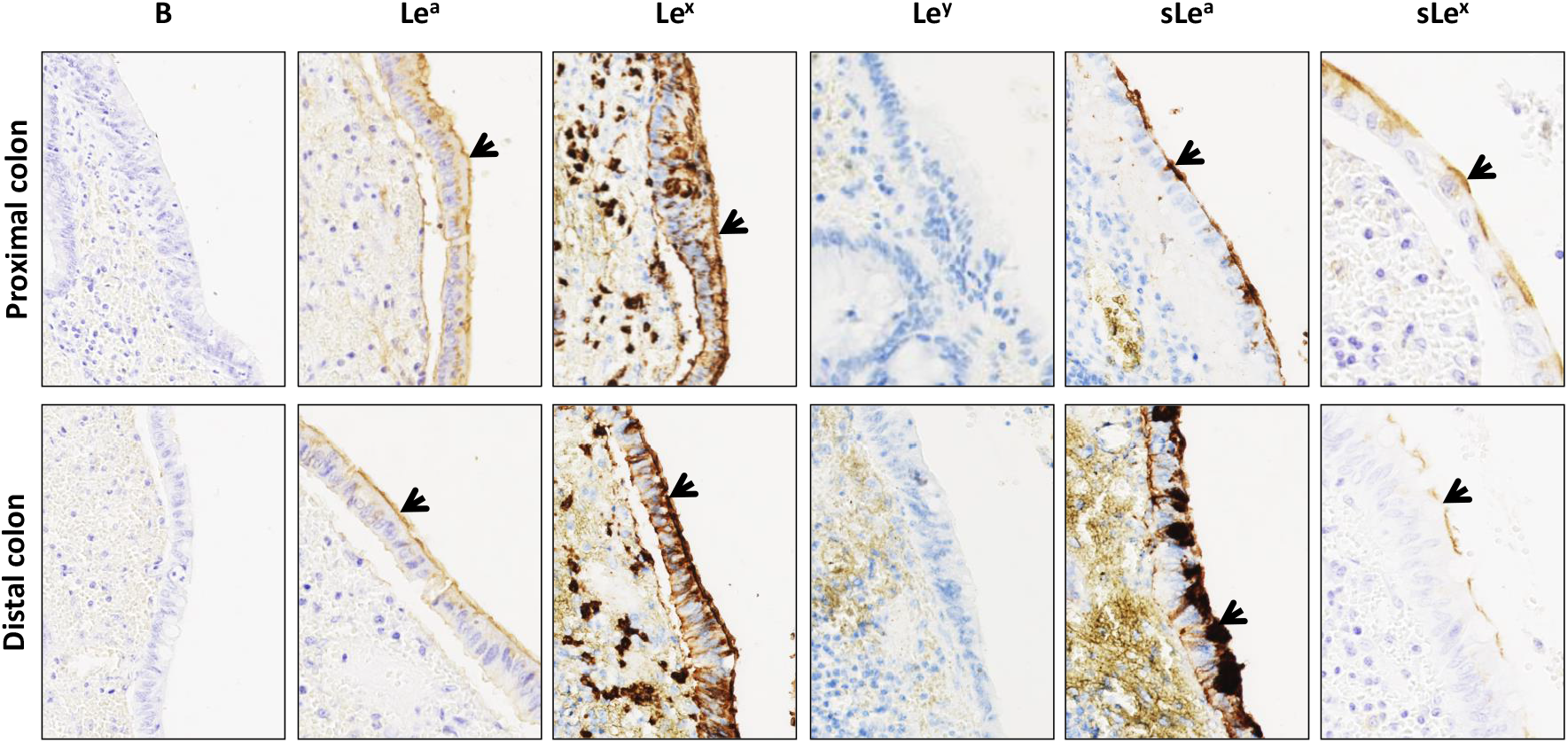
Blood group B, Lewis antigens and sialylated Lewis antigens detection in regenerative mucosa of samples of proximal colon and distal colon from non-secretor Crohn’s disease patient 14NS. The anatomical site of each sample is indicated on the left side of the panel. The detected HBGA are indicated on top of each row. Positive HBGA detection is shown by light or dark brown staining and with arrows. The panels are shown at magnification x400.

### Characterization of GII.4 and GII.17 attachment to non-secretor regenerative mucosa

Our next step was to identify the ligands involved in GII.4 and GII.17 HuNoV interaction in regenerative mucosa. In a previous report, we showed that Le^a^ and to a lesser extent Le^x^ antigens were involved in GII.4 attachment to regenerative mucosa (16). We subsequently hypothesized that the Le^a^ antigen would also be involved in the attachment of GII.4 and GII.17 HuNoV VLPs to non-secretor colonic regenerative mucosa. Le^a^-specific mAbs were used for the competition experiments while Le^b^-specific mAbs were used as a negative control. The binding of GII.4 and GII.17 HuNoV VLP was abolished in regenerative mucosa samples from the proximal and distal colon when tissue slides were preincubated with Le^a^-specific mAbs, while Le^x^-specific mAbs failed to inhibit VLP binding (Figure 10). Our observations confirmed that the Le^a^ antigen was the main ligand for HuNoV VLP interaction in inflammatory tissues sampled from a non-secretor.

**Figure 10:**
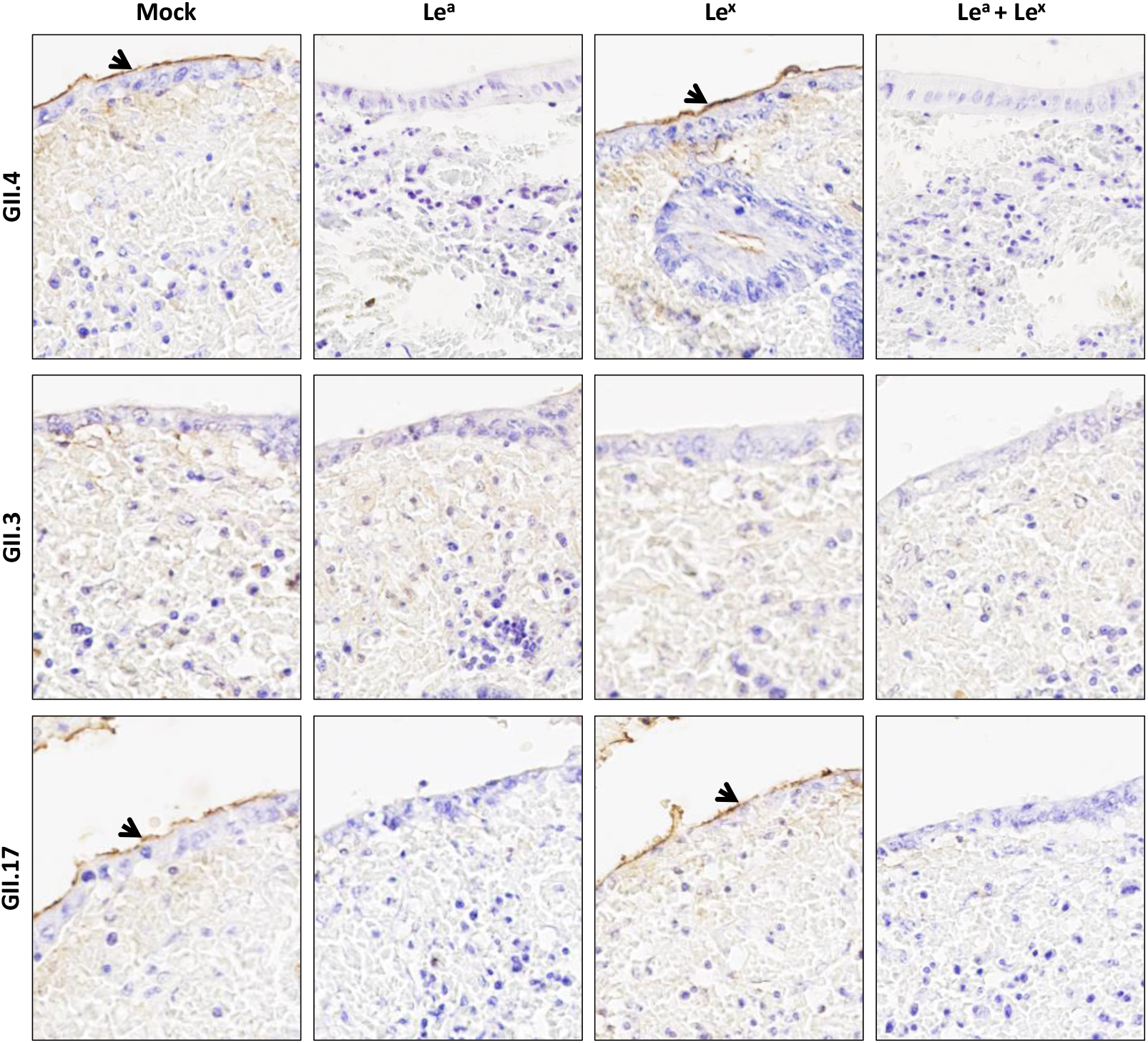
VLP competition experiments performed on regenerative mucosa in samples of proximal colon and distal colon from non-secretor Crohn’s disease patient 14NS (magnification x400). Preincubation of mAbs and mock control are indicated on top of each row. The following incubation of VLP is indicated at the left of each panel. Positive VLP detection is indicated by brown staining and with arrows.

## DISCUSSION

Human norovirus infection is strongly correlated to the expression of HBGAs due to the active FUT2 gene in the small intestine, as recently exemplified in a recent study of a large pediatric cohort (38). Nevertheless, there is a paucity of data concerning HuNoV infection in non-secretor individuals. Since the identification of HBGAs as ligands of HuNoVs, the binding profiles of HuNoVs have mostly been characterized using synthetic carbohydrates and saliva samples with bacterially-expressed P particles and baculovirus-expressed VLPs (39-41). Most HuNoV genotypes are effective binders of secretor saliva containing ABO(H) antigens, but this is not the case for saliva from non-secretor individuals (36, 42). Previous studies showed that the GII.4 2002a variant weakly bound non-secretor saliva (43). We also showed that 2006b/Den Haag variant and 2007/Osaka variant were efficient binders of non-secretor saliva, whereas other GII.4 variants were not (36). Here, we used a panel of non-secretor saliva to assess and confirm the binding capacity of three variants: GII.4 Osaka, GII.17 Kawasaki 308 and GII.3. GII.4 Osaka. The competition experiments demonstrated the pivotal role of the Le^a^ antigen in the binding of GII.4 VLPs in saliva samples. These experiments also showed an absence of GII.3 binding on saliva, confirming the results of our previous study (37). In this previous study, we produced GII.3 VLPs derived from a strain (SW4) detected in a non-secretor individual. This GII.3 VLP bound none of the non-secretor saliva samples including saliva from the individual where it originated. Our results were thus surprising because it could be assumed that the HuNoVs found in non-secretor individuals would bind the saliva from those individuals provided that HBGAs are involved into the recognition of HuNoVs. Our results were inconsistent with epidemiological observations, but we hypothesized that non-HBGA ligands played a role in the recognition of the GII.3 HuNoV. One could also consider that the levels of expression of HBGA in saliva might not entirely reflect the physiological expression of HBGA at the intestinal level. The lower abundance of Le^a^ in saliva and three-dimensional conformation of this antigen may be responsible for the poor recognition of the viral particles and explain the absence of binding of HuNoV on the saliva of non-secretor individuals while it still manages to infect the intestinal cells. To investigate this hypothesis, our novel study analyzed HuNoV interactions on duodenal epithelium samples from non-secretor individuals. We observed specific binding of VLPs from the three genotypes for the majority of the non-secretor duodenum tissues. Therefore, our data confirmed that GII.3 and GII.17 HuNoV VLPs may interact with samples of intestinal tissues but not with the corresponding saliva samples. The Le^a^ antigen was the main ligand involved in the recognition of the genotypes GII.3, GII.4 and GII.17 in non-secretor individuals. A HIE-based study previously showed that GII.3, GII.4 and GII.17 recognize ligands at the surface of non-secretor enteroids but with a lower binding amplitude than that observed for secretor enteroids (9). Differences in glycan conformation, structure and abundance might also explain the variations in HuNoV VLP binding. In the digestive tract, the proximal and distal colon exhibit a pan-mucosal expression of the Le^a^ antigen, alone or in its monosialylated or disialylated forms, without showing HuNoV binding (44-46). The expression level and thus the abundance of the Le^a^ antigen could differ between the colonic and ileal mucosa, which could partly explain why GII.17 and GII.3 interact with healthy duodenum but not with distal and proximal colon, where the expression levels of Le^a^ ligands are substantially lower. Variations in the levels of Le^a^ expression could therefore explain our findings for samples 10NS through 13NS. It is also conceivable that other ligands associated with Le^a^ are involved in HuNoV recognition and might be essential for HuNoV replication. This is the case for murine norovirus, in which the CD300lf protein is the viral receptor whereas ganglioside GD1a might be considered as a co-receptor (47, 48). These observations suggest the need for future research on the role of non-HBGA ligands during HuNoV infection.

The Le^a^-driven binding of GII.3 VLPs is coherent with previous epidemiological results that documented GII.3-related gastroenteritis in non-secretor individuals (37). Inversely, GII.4 and GII.17 outbreaks are strongly associated with secretor-positive status (49). Although rare, GII.4 infections of non-secretor patient bearing nonsense mutation G428A have been documented and support the concept of abortive or minor infections by the GII.4 HuNoVs (50-53). Of note, GII.4 infections have also been reported in “weak” secretors with the A385T missense mutation (54, 55). Similar epidemiological observations have been made in studies of GII.17 HuNoVs, in which approximately 8% of non-secretor individuals were infected (56). A retrospective serological analysis using GII.4 VLPs showed that non-secretor individuals had developed GII.4-specific serological response (57). For non-secretor patients, GII.4-positive serology might reflect an aborted GII.4 HuNoV infection, which may depend on other factors (i.e., age, health status, immunity, microbiota) (58). It is assumed that the odds of HuNoV exposure is the same for secretor and non-secretor individuals. For non-secretor individuals, it can be hypothesized that HuNoV binding occurs through the recognition of Le^a^ antigen, and is not necessarily followed by HuNoV internalization and replication, especially for GII.4. Studies conducted in various Asian, American, European and African populations have already revealed significant polymorphisms in the *FUT3* gene, which is involved in the complex pathways of Lewis antigen synthesis in erythrocytes and secretions (59-63). As mentioned above, the differential levels of expression of the *FUT3*-driven Lewis antigens (especially Le^a^ and Le^x^) might explain the variations in HuNoV binding patterns in the intestinal tissues of non-secretor individuals. Detailed analysis of HBGA expression and *FUT3* polymorphisms might provide information about the capacity of HuNoV to bind to the intestinal tissues of non-secretor individuals. The use of mutant enteroids for *FUT2* and *FUT3* could also provide new information regarding HBGA expression and HuNoV interactions in secretors and non-secretors, as described recently (9).

Recent developments in the culture of HuNoV using HIE have provided groundbreaking new tools for the study of the molecular mechanisms driving HuNoV infections. GII.3, GII.4 and GII.17 replication has been studied on non-secretor and secretor HIEs with a knocked out FUT2 gene (9). These experiments showed that GII.4 and GII.17 replication was almost entirely restricted to secretor HIE with an active FUT2 gene (5, 9). However, a weak replication was observed for GII.17 in non-secretor HIEs. Surprisingly, only GII.3 and not GII.17 could efficiently replicate in knock-out HIE for the FUT2 gene. It is probably overly simplistic to assume that Le^a^ is the sole ligand recognizing HuNoV in non-secretor individuals, and HuNoV replication is likely conditioned by other factors.

Nevertheless, there are some limitations concerning the use of enteroids to study HBGA expression, even when using identical tissue-derived enteroids. Enteroids are generally derived from intestinal stem cells, located at the base of the mucosa, thus possessing a greater capacity for proliferation. It is also worth mentioning that they are well-known for being the cells of origin of intestinal neoplasia (64). Therefore, enteroids do not necessarily reflect physiological or inflammatory conditions in the intestine. Indeed, the higher proliferation capacity of intestinal cryptic niches is often correlated to their greater pluripotency, for instance the overexpression of the HBGAs such as sLe^a^, sLe^x^ and Le^y^ in the small and large intestines (46). Similarly, digestive neoplasms show greater expression of Lewis antigens, providing a greater capacity for proliferation and dissemination (65-67). A study conducted by Zheng et al. demonstrates the high expression of type 2 blood group antigens such as Le^x^ and sLe^x^ antigens in enteroids, regardless of the secretor status (68). Future studies using both histological assays and HIE replication assays could help decipher the norovirus susceptibility and replication capacity of different intestinal cell subtypes. Finally, other relevant factors such as bile acid secretion in the intestinal lumen, the composition of the intestinal barrier, and gut microbiota interactions with noroviruses should be taken into account in future studies focusing on HuNoV pathogenicity.

In our previous study, the putative role played by Le^a^, and to a lesser extent by Le^x^, was demonstrated through HuNoV recognition on inflammatory and regenerative colonic mucosa in secretor IBD patients. Surprisingly, VLP attachment was genotype-dependent in the regenerative mucosa, whereas only GII.4 and GII.17 VLPs interacted with mucosae from the proximal and distal colon. For both quiescent and regenerative colonic mucosae in IBD patients, VLP binding suggested that Lewis antigens (mainly Le^a^ and sLe^a^) were involved in HuNoV binding in the absence of ABH(O) antigens in both secretor-dependent tissues (i.e. small intestine and proximal colon) and secretor-independent tissues (i.e. distal colon). As for inflammatory and regenerative tissues, mouse models showed that α1,3-fucosyltransferase and α2,3 sialyltransferase 4 activity is increased during colitis, which might explain increased expression of sialylated lewis antigens and thus increased HuNoV binding (69, 70).

The major drawback of our study is that the analyses are based on tissue samples from a single patient. Future research will obviously require the observation of several Crohn’s disease cases given that the likelihood of finding non-secretor Crohn’s disease patients with flare-ups is low. Furthermore, the use of intestinal biopsies derived from non-secretor individuals remains challenging due to the small amount of tissue in biopsy samples, the difficulty of *FUT2* screening in formalin-fixed paraffin-embedded (FFPE) tissue samples, and bioethics regulations. In future studies, data obtained with VLPs should be supported using native HuNoV in HIE systems, as described previously for the study of neutralizing mAbs (71). That being said, because the use of HIE is technically challenging, time-consuming and costly, the study of HuNoV attachment using VLPs and histological tissues provides a good alternative in the absence of cultivated organoids. The strategy used for the study of HuNoV replication in immunocompromised patients paves the way for new research in IBD patients suffering from HuNoV infection (8). Similar studies combining epidemiological data and immunohistological analyses should be undertaken in IBD patients. The physiological consequence of HuNoV binding to Le^a^ should also be clarified from an immunological point of view.

In summary, epidemiological surveys and HIE-based studies have shown that secretor individuals are far more susceptible to HuNoV infections. However, similar exposure to HuNoVs can also cause gastroenteritis in non-secretor individuals in rare circumstances. Our observations suggest that, in healthy non-secretor individuals, the attachment of viral particles occurs via the Le^a^ antigen in the duodenum. Additionally, the Le^a^ antigen could play a pivotal role in inflammatory mucosae from the small intestine and the colon, irrespective of secretor status. Future work in the context of IBD should address whether HuNoV infection is a cause or a consequence of inflammation. Such studies should obviously be extended to other enteric viruses that interact with HBGAs, such as the rotavirus (72-74). Finally, future research is warranted to elucidate whether viral infections could be triggering or aggravating factors in other intestinal conditions such as celiac disease or irritable bowel syndrome.

## MATERIALS AND METHODS

### Screening and selection of tissue samples

Formalin-fixed paraffin-embedded (FFPE) tissue samples were selected from the archives of the Pathology Department of the Dijon University Hospital. Approval for the study (reference 18.11.29.52329) was granted by the French national ethics committee (CPP19002). One hundred and twenty-one duodenal biopsy samples were selected and read by two pathologists (L.M., G.T.) to rule out inflammation and dysplasia. We also retrieved samples of healthy tissue derived from twenty-five surgical resection specimens (16 Crohn’s disease and 9 ulcerative colitis patients).

Screening for the *FUT2* gene and HBGA detection was performed in all samples devoid of lesions. Thirty-four saliva samples from non-secretor individuals from French and Tunisian cohorts were used for salivary binding assays (36, 37) (37, 38). Their use was previously approved by the Nantes University Hospital Review Board for the French cohort (study no. BRD02/2-P). For the Tunisian cohort, the study was approved by the Ethics Committee of the Fattouma Bourguiba Public Hospital at Monastir (Tunisia) (committee decision of the 9th of May 2013), and informed consent was obtained from the parents of the involved children.

### FUT2 genotyping

Duodenal biopsies and IBD tissue samples were screened for the G428A mutation characterizing the non-secretor genotype of the FUT2 gene. DNA extraction from FFPE tissues and FUT2 genotyping by sequencing were described previously (15, 75).

### Virus-like particles and antibodies

For this study, we used Virus-Like Particles (VLP) derived from HuNoV strains, incubated on saliva and tissue samples. Cairo-4 GII.4 VLP were derived from an Osaka/2007 variant (36, 76). SW4 GII.3 VLP originated from a non-secretor child suffering from viral gastroenteritis (37). The E12905 isolate is a Kawasaki308 (Kawa308) variant GII.17 from an 86-year-old patient. The Kawa308 variant was first described in Hong Kong (77). Cloning details and production of recombinant baculovirus is available upon request. Production and cesium gradient purification of the baculovirus-expressed VLPs were previously described (36, 78). GII.4 VLP were detected with in-house specific mAbs recognizing conformational epitope and labeled with horseradish peroxidase (HRP). GII.3 and GII.17 VLPs were detected with rabbit specific polyclonal sera (bioMérieux, Marcy l’Etoile, France). The Le^x^, Le^a^, Le^b^, sLe^a^ and sLe^x^ antigens were detected with mAbs MEM-158 (Sigma-Aldrich France), 7-Le (Sigma-Aldrich France), 2-25LE, NS-1116-19.9 (Dako, USA) and CSLEX-1 (Becton-Dickinson, USA), respectively.

### Saliva binding assay and competition experiments

For the salivary VLP binding assays, an Enzyme-Linked Immunosorbent Assay (ELISA) was performed as described previously, except that 1000-fold diluted saliva and 500 ng of purified VLP were used per assay (10). Salivary binding patterns of HuNoV VLP were previously described (36). Briefly, GII.4, GII.3 and GII.17-bound VLPs were detected by 5000-fold diluted mAbs, 2000-fold diluted and 10.000-fold polyclonal sera, respectively. All primary antibodies were diluted in phosphate-buffered saline (PBS) with 4% dry milk (blotto) and incubated for 1 h at 37°C. For the detection of the polyclonal sera, peroxidase-conjugated anti–rabbit antibody (Sigma, France) was incubated for 1 h at 37°C. Peroxidase activity was detected with 3, 3’, 5, 5’-tetramethyl benzidine (TMB from KPL/Eurobio, Courtaboeuf, France). The reaction was stopped after 10 min of incubation at room temperature with 2N HCl prior to absorbance reading at 450 nm. The background was arbitrarily fixed at 0.2 OD (optical density).

Four non-secretor saliva samples were used for the competition experiments using Cairo 4 GII.4 VLP. Because the HBGA content is time- and donor-dependent, preliminary binding assays with Cairo 4 VLPs were conducted to determine the dilutions that resulted in binding values ranging between 0.5 and 1.0 OD450. Salivary samples were diluted either 1000-, 8000- or 16.000-fold in carbonate/bicarbonate buffer and coated at 37°C, overnight. The plates were washed 4 times with PBS and incubated with 2-fold serial dilution starting at 500 ng/well down to 8 ng/well of either Le^a^-, Le^b^- or Le^x^-specific mAbs diluted in PBS and incubated overnight at 37°C. The ELISA plate was then quenched with PBS-4% blotto for 1.5 h at 37°C before incubating 250 ng/well of GII.4 purified VLP for 2 h at 37°C. Bound VLPs were then detected with GII.4-specific HRP-labeled mAbs, as described above, and HRP activity was detected with TMB incubated for 10 min at room temperature.

### Histological analysis and immunodetection on tissue samples

Four µm-thick microtome sections were cut from FFPE samples. Sections were deparaffinized, quenched and pretreated for endogenous peroxidase inhibition, as previously described (15). Five µg/ml of in-house purified GII.4, GII.3 or GII.17 VLP diluted in PBS with BSA 1% were incubated on each slide overnight at 4°C. The 5B9 HRP-labeled antibodies and rabbit anti-immunoglobulin HRP-labeled antibodies were all detected using 3’-3’-Diaminobenzidine for 1.5 min at room temperature, and counterstained with hematoxylin (Dako, Agilent Technologies, USA), as previously described (15). Le^a^ and Le^x^ were detected with 0.5 µg/ml of mAbs MEM-158 and 7Le (*Sigma*-Aldrich, Germany), respectively. Sialyl-Lewis a (sLe^a^) and Sialyl-Lewis x (sLe^x^/CD15s) were detected with 2 µg/ml of mAbs NS-1116-19.9 (Dako, Agilent Technologies, USA) and CSLEX-1 (Becton Dickinson, USA), respectively. Antibody incubation and detection was performed on a Dako^®^ OMNIS^®^ automaton (Dako, Agilent Technologies, USA).

### Competition experiments on tissue samples

Competition experiments using VLP were performed on duodenal biopsies and IBD tissues derived from non-secretor individuals. Four µm-thick sections were obtained from duodenal biopsies and macrodissected blocks derived from surgical resection specimen samples. Slides were preincubated with 25 µg of mouse mAbs directed against Le^a^ (clone 7LE, Thermo-Fischer), Le^b^ (clone 2-25LE, Thermo-Fischer) or Le^x^ (clone MEM-158), combined or alone, prior to VLP incubation, as described above.

## ACKNOWLEDGMENTS

This work was supported by the National Reference Center for Viral Gastroenteritis and the Dijon-Bourgogne University Hospital (France). Georges Tarris received a fellowship from the Faculty of Medicine of the University of Burgundy (Dijon, France). We would like to thank Suzanne Rankin for providing editorial assistance.

## DATA AVAILABILITY STATEMENTS

Digitized images (WSI format) from the histological analyses and HuNoV VLPs used in the manuscript are available upon request.

